# Reconstitution of ribosome self-replication outside a living cell

**DOI:** 10.1101/2022.08.29.505692

**Authors:** Yuishin Kosaka, Yumi Miyawaki, Megumi Mori, Shunsuke Aburaya, Mao Fukuyama, Mitsuyoshi Ueda, Wataru Aoki

**Author notes:** Correspondence should be addressed to Wataru Aoki.

## Abstract

Ribosome biogenesis, a recursive process of pre-existing ribosomes self-replicating nascent ones, is pivotal in the self-replication of life. In *Escherichia coli*, three ribosomal RNAs (rRNAs) are transcribed, and 54 ribosomal proteins (r-proteins) are synthesized by pre-existing ribosomes as structural components^1, 2^. They are cotranscriptionally assembled in a cooperative hierarchy under the support of ∼100 accessory factors^1–3^. The reconstitution of ribosome biogenesis outside a living cell is an essential goal to understand the self-replication of life. However, this goal could not have been achieved so far due to its complexity. Here, we report the successful *in vitro* reconstitution of the entire ribosome biogenesis process. We hypothesized that mimicking *in vivo* ribosome biogenesis^1–6^ could result in *in vitro* ribosome biogenesis. Specifically, we found that coactivating the transcription of an rRNA operon, as well as the transcription and translation of 54 r-protein genes encoding r-proteins, and the coordinated ribosomal assembly in a cytoplasm-mimicking reaction solution, resulted in highly efficient *in vitro* reconstitution of ribosome biogenesis. Our achievement represents a critical step toward revealing fundamental principles underlying the self-replication of life and creating self-replicating artificial cells^7^. We also succeeded in engineering rRNA and r-proteins by only adding mutant ribosomal genes in the reaction, enabling high-throughput and unconstrained creation of artificial ribosomes with altered or enhanced functionality^8–12^.

## Main

Ribosome biogenesis is a recursive process in which ribosomes, the universal decoders of the genetic code, are self-replicated by pre-existing ribosomes. This process is pivotal in the self-replication of life and is universally conserved across living organisms. In *Escherichia coli*, three rRNAs (16S, 23S, and 5S) are transcribed by the RNA polymerase enzyme and 54 r-proteins are synthesized by pre-existing ribosomes as structural components^1, 2^. They are cotranscriptionally assembled in a cooperative hierarchy through multiple parallel assembly pathways^13–22^. The assembly process is supported, modified, and modulated by ∼100 accessory factors^1, 2^. All these steps concurrently occur in the cytoplasmic space in a highly coordinated manner, resulting in the synthesis of the 2.5-MDa 70S ribosome, consisting of the 30S small and 50S large subunits (SSU and LSU, respectively), in a few minutes^23^. The SSU and LSU are essential for translation and contain the decoding and the peptidyl transferase center, respectively. Ribosomes play multifaceted roles in healthy cells, and ribosome biogenesis dysregulation leads to the development of various aberrant states such as cell death and cancer^24^.

Reconstituting ribosome biogenesis outside a living cell is an essential goal in biology to understand the self-replication of life. Intensive scientific efforts have been invested in achieving this goal for decades. Ribosome assembly mapping revealed assembly order and intermediates, as well as thermodynamic and kinetic parameters^25–27^. The *in vitro* integrated synthesis, assembly, and translation (iSAT) realized the coupling of rRNA synthesis and ribosome assembly using purified r-proteins^4, 5, 28, 29^. These efforts in nonautonomous ribosome assembly with purified r-proteins encouraged attempts to reconstitute ribosome biogenesis *in vitro*. A study describes an attempt to cogenerate r-proteins from DNA templates in an *in vitro* one-pot reaction^30^. Another study conducted simultaneous expression of SSU structural components and certain accessory factors on a chip in an attempt to reconstitute SSU biogenesis^31^. The latter study reproduced hallmarks of SSU biogenesis on a chip; however, nascent SSU activity as the decoding center was not confirmed^31^. To the best of our knowledge, reconstitution of LSU biogenesis, a far more complex process than SSU biogenesis^2^, has not even been attempted yet. Hence, a big leap needs to be made forward to reconstitute ribosome biogenesis *in vitro*.

In this study, we report the first successful *in vitro* reconstitution of the entire ribosome biogenesis process in *E. coli*. We hypothesized that mimicking the *in vivo* ribosome biogenesis process^1, 2, 4, 5^ and cytoplasmic chemical conditions^4, 6^ could result in *in vitro* ribosome biogenesis. Specifically, our approach involved coactivating the transcription of an operon encoding three rRNAs, the transcription and translation of 54 genes encoding r-proteins, and the coordinated assembly of ribosomes in an optimized *E. coli* S150 cell extract, containing the ∼100 accessory factors for ribosome biogenesis and imitating cytoplasmic chemical conditions. To test our hypothesis, we developed a highly specific, sensitive reporter assay to detect the translational activity of nascent ribosomes. The reporter assay allowed for the stepwise, combinatorial exploration of the reaction conditions, leading us to successful reconstitution of the entire ribosome biogenesis process *in vitro*, that is, autonomous self-replication of the 2.5-MDa 70S ribosome by concurrent transcription, translation, processing, modification, modulation, and assembly in a single reaction space. The reconstituted *in vitro* ribosome biogenesis allows us for more freedom in controlling the process of ribosome biogenesis. Therefore, this achievement would generate a widespread impact on understanding the self-replication of life, elucidating the ribosome assembly process^1, 2^, revealing the multifaceted roles of ribosome biogenesis in cell physiology^24^, creating self-replicating artificial cells^7^, and designing artificial ribosomes with altered or enhanced functionalities^8–12^.

### Development of a highly specific reporter assay for nascent ribosome detection

In an attempt to reconstitute ribosome biogenesis *in vitro*, a highly specific reporter assay for detection of the nascent ribosome translational activity would be required as pre-existing and nascent ribosomes would coexist in a single reaction space. Translational efficiency is mainly determined by the RNA–RNA base pairing between the Shine–Dalgarno (SD) and anti-Shine–Dalgarno (ASD) sequences of mRNA and the 16S rRNA, respectively^32^. Therefore, generation of new SD and ASD leads to the development of orthogonal translation systems^33–36^ useful for detecting nascent artificial ribosomes (**Fig. 1a**). Among them, a two-sided orthogonal translation system would be superior in specificity and sensitivity. A previous study described that certain pairs of orthogonal SDs and ASDs (oSDs and oASDs, respectively) exhibit two-sided orthogonality in *E. coli*^34^. However, whether any oSD·oASD pairs exhibit two-sided orthogonality *in vitro* remains elusive^28^.

**Fig. 1.**
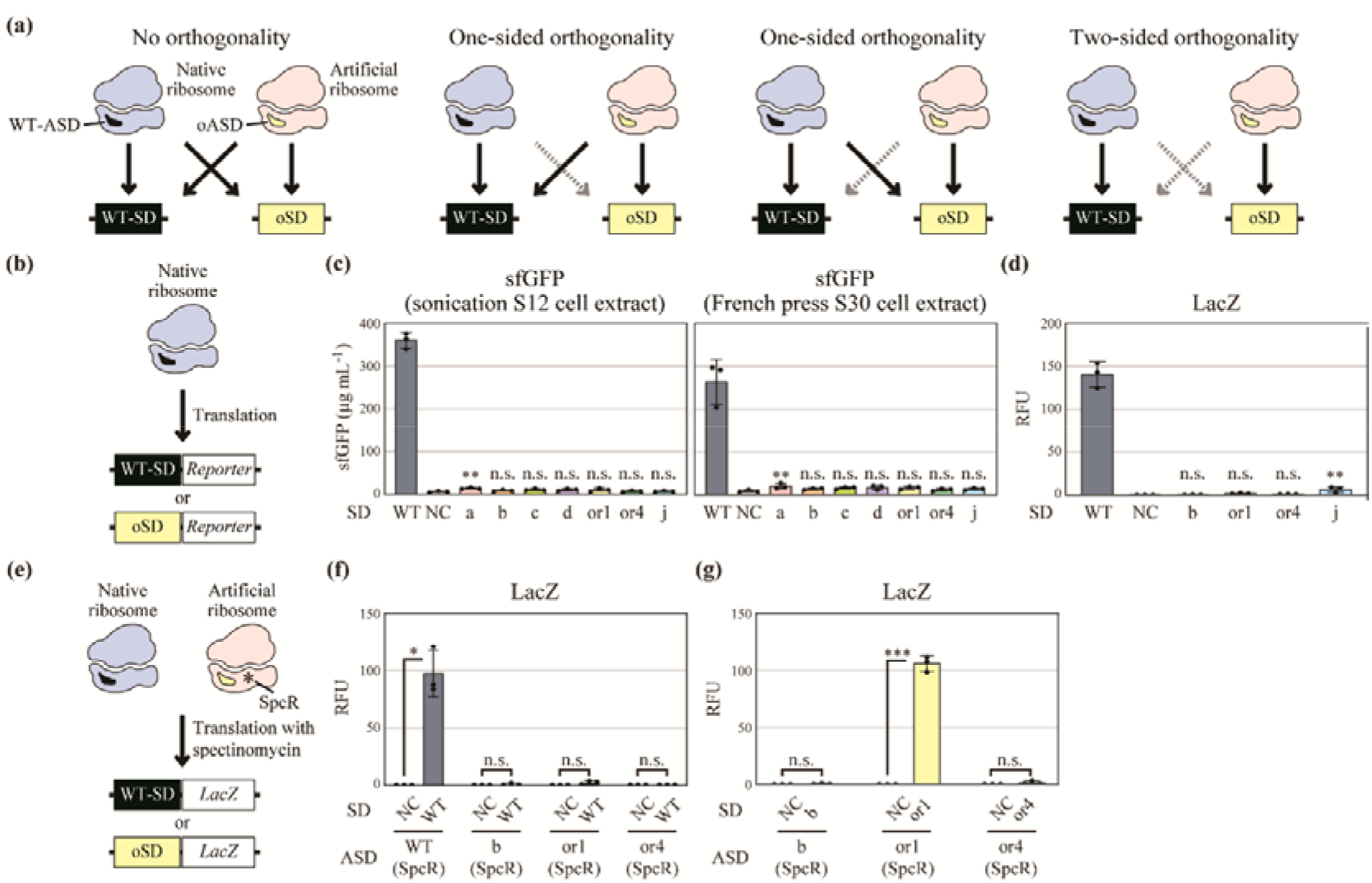
Screening orthogonal oSD·oASD pairs with two-sided orthogonality *in vitro*. **(a)** Four types of orthogonalities. Black and gray arrows indicate functional and nonfunctional interactions, respectively. SD, Shine–Dalgarno sequence; ASD, anti-Shine–Dalgarno sequence; oSD, orthogonal SD; oASD, orthogonal ASD. **(b)** Experimental scheme to screen oSDs that do not interact with native ribosomes in cell extracts. **(c)** oSD selection. Either a WT-SD–sfGFP or an oSD–sfGFP reporter (named a, b, c, d, or1, or4, and j) was mixed with S12 or S30 cell extracts prepared using BL21 Star^TM^ (DE3) *lacZ::kmr*. NC, negative control without a reporter. The data represent the mean ± SD (n = 3). **, *p* < 0.01; n.s., not significant; Dunnett’s test against NC. **(d)** Further oSD selection. Either a WT-SD–LacZ or an oSD–LacZ reporter (b, or1, or4, and j) was mixed with S12 cell extracts. RFU, relative fluorescence unit; NC, negative control without a reporter. The data represent the mean ± SD (n = 3). **, *p* < 0.01; Dunnett’s test against NC. **(e)** Experimental scheme to screen oSD·oASD pairs with two-sided orthogonality in cell extracts. Cell extracts were prepared using BL21 Star^TM^ (DE3) *lacZ::frt* expressing an artificial rRNA operon with WT-ASD or oASD (b, or1, or or4) and C1192U spectinomycin resistance (SpcR). **(f)** Screening oASDs that do not interact with the WT-SD–LacZ reporter. The cell extracts were mixed with the WT-SD–LacZ reporter and spectinomycin. NC, negative control without the reporter. The data represent the mean ± SD (n = 3). *, *p* < 0.05; Welch’s *t*-test. **(g)** Screening oSD·oASD pairs with two-sided orthogonality. The cell extract was mixed with the cognate oSD–LacZ reporter and spectinomycin. NC, negative control without a reporter. The data represent the mean ± SD (n = 3). ***, *p* < 0.001; Welch’s *t*-test.

We selected seven oSD·oASD pairs^34–36^ (named a, b, c, d, or1, or4, and j) as candidates to screen two-sided orthogonal translation systems available in *E. coli* cell extracts **(Supplementary Information 1)**. First, we designed an experimental scheme to select oSDs that do not interact with native ribosomes in the cell extracts and designed fluorescent reporter constructs for each member of the selected candidate pairs (**Fig. 1b**). Either a WT-SD–sfGFP or an oSD–sfGFP reporter was mixed with two types of cell extracts (sonicated S12 or French press S30) containing native ribosomes. We observed that six oSDs (b, c, d, or1, or4, and j) did not show any functional interaction with the native ribosomes (**Fig. 1c**). Both cell extracts showed similar profiles; hence, we used the S12 extracts for the following screening processes due to their ease of preparation. We thus further investigated the orthogonality of the top four oSDs (b, or1, or4, and j) using LacZ reporters that were more sensitive than the GFP reporters, and discovered that three oSDs (b, or1, and or4) displayed strong orthogonality against the native ribosomes (**Fig. 1d**).

Next, we designed an experimental scheme to screen oSD·oASD pairs with two-sided orthogonality in cell extracts (**Fig. 1e**). We prepared functional cell extracts using *E. coli* expressing an artificial rRNA operon with WT-ASD or oASD (b, or1, or or4) and C1192U spectinomycin resistance (SpcR)^37^ in the 16S rRNA **(Extended Data Fig. 1a)**. The cell extracts containing artificial ribosomes with b-, or1-, or or4-oASD did not generate reporter signals when mixed with the WT-SD–LacZ reporter and spectinomycin (**Fig. 1f** **and Extended Data Fig. 1b)**. When mixed with the cognate oSD–LacZ reporter and spectinomycin, the cell extract containing artificial ribosomes with or1-oASD generated a strong reporter signal (**Fig. 1g**). We verified in a follow-up control experiment that the reporter signal certainly derived from the or1-oSD·oASD pairing **(Extended Data Fig. 1c)**. These results showed that the or1-oSD·oASD pair exhibited strong two-sided orthogonality in the cell extracts. We have not pursued the reason why b- and or4-oSD·oASD pairs were nonfunctional (**Fig. 1g**). A potential explanation is that our expression vectors did not allow the expression of the artificial rRNAs with b- or or1-oASD in *E. coli*.

Encouraged by the success to develop the highly specific reporter assay, we conducted a preliminary trial to reconstitute SSU biogenesis *in vitro*. We coactivated the transcription of the artificial rRNA operon with or1-oASD and SpcR, the transcription and translation of 21 SSU r-protein genes, and the coordinated assembly in an optimized S150 cell extract, containing the ∼100 accessory factors for ribosome biogenesis and imitating cytoplasmic chemical conditions. However, we observed no nascent SSU-derived reporter signal **(Extended Data Fig. 2)**, confirming the difficulty to reconstitute such a complex process *in vitro*. Therefore, we conceived that a highly sensitive assay would also be required to detect the translational activity of nascent ribosomes to explore the reaction conditions which would enable ribosome biogenesis *in vitro*.

### Highly sensitive detection of the artificial ribosome translational activity

We devised a deep-leaning-assisted automated femtoliter droplet assay for sensitive, scalable, and objective detection of the artificial ribosome translational activity. A femtoliter droplet assay, in which a tiny amount of the reaction solution is confined to femtoliter droplets, allows for highly sensitive enzymatic activity detection^38^ (**Fig. 2a**). However, scalable and objective femtoliter droplet assay analysis is generally difficult as the bright-field images of the droplets have an extremely low contrast, hampering precise droplet segmentation. To address this problem, we developed a deep-learning-assisted automated analysis pipeline (**Fig. 2b**), using a trained U-Net^39^ deep-learning model to transform bright-field droplet images into binary segmented images (droplet or background) with >90 % accuracy **(Extended Data Fig. 3)**. We used the binary segmented images to extract area, fluorescence intensity, and other features of each droplet by automated particle analysis using ImageJ^40^.

**Fig. 2.**
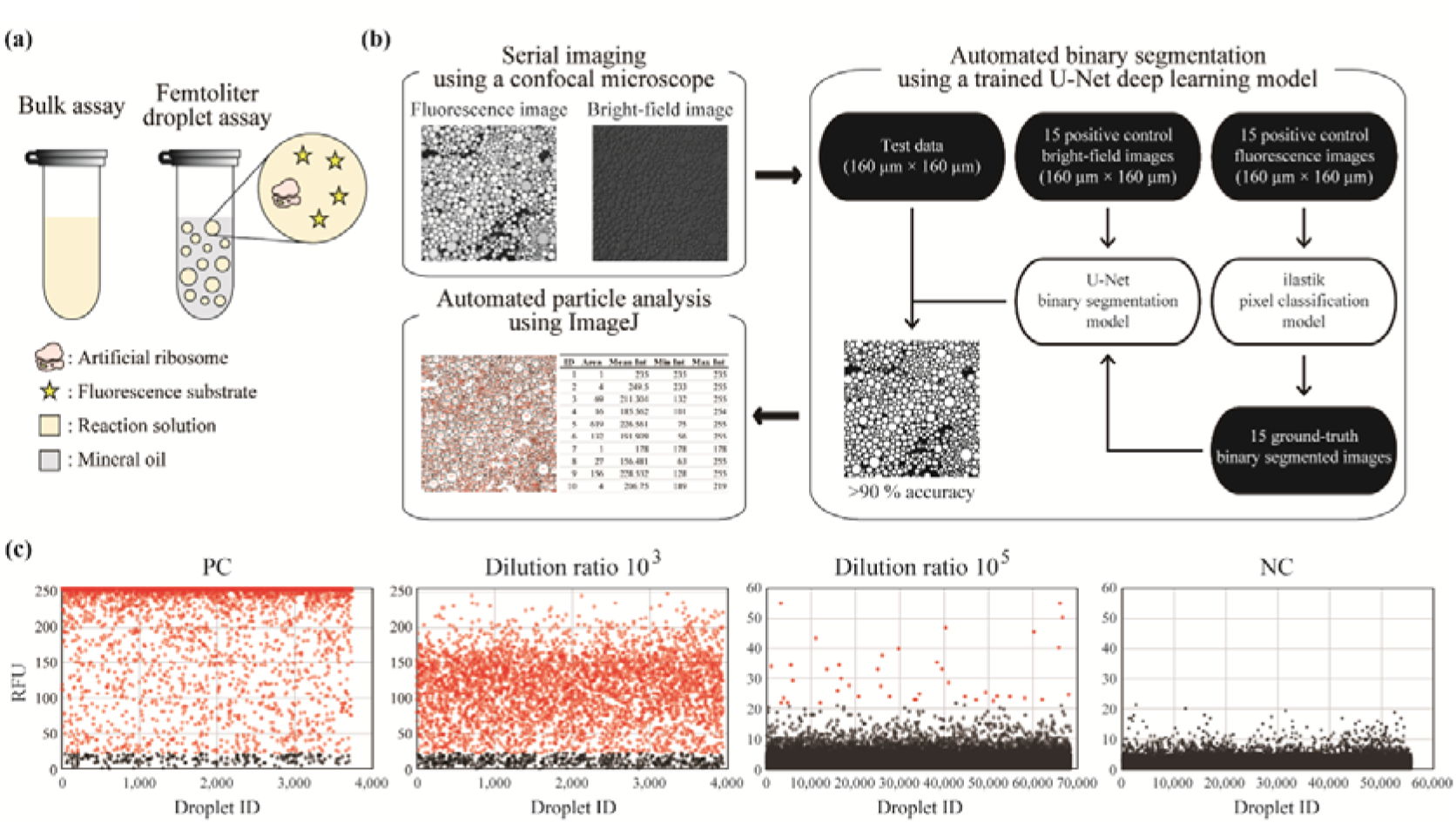
Highly sensitive detection of the artificial ribosome translational activity. **(a)** Comparison between a conventional bulk assay and a femtoliter droplet assay. **(b)** Deep-learning-assisted automated femtoliter droplet assay. After cell-free transcription and translation in femtoliter droplets, bright-field and fluorescence images of the droplets are obtained using a confocal fluorescence microscope. The bright-field images are binary segmented (droplet or background) using a trained U-Net deep-learning model. The binary segmented images are obtained for particle analysis using ImageJ, and the results are redirected to corresponding fluorescence images. Finally, we obtain the features of each droplet including area and relative fluorescence unit (RFU). **(c)** Highly sensitive detection of the artificial ribosome translational activity. We prepared two types of S12 cell extracts; one contained native ribosomes and 4.9 μM of artificial ribosomes with or1-oASD and C1192U spectinomycin resistance (SpcR) and the other only native ribosomes. The cell extract containing the artificial ribosomes was diluted by the control cell extract at the indicated ratio. The cell-free transcription and translation in femtoliter droplets were carried out in the presence of the or1-oSD–LacZ reporter and 100 μM of spectinomycin. In the scatter plots, the vertical and the horizontal axes indicate the mean RFU and the ID of each droplet, respectively. Droplets over the threshold (the mean RFU ≥ 22) are indicated in red. PC, positive control without dilution; NC, negative control using only the control cell extract.

Next, we evaluated the sensitivity of the deep-learning-assisted automated femtoliter droplet assay. We prepared two types of S12 cell extracts: one contained native ribosomes and 4.9 μM of artificial ribosomes with or1-oASD and SpcR **(Extended Data Fig. 4)**, and the other was a control cell extract containing only native ribosomes. We performed a control experiment by mixing the control cell extract with the or1-oSD–LacZ reporter. Unexpectedly, we observed native ribosome-derived fluorescence **(Extended Data Fig. 5a),** not detected in the bulk assay (**Fig. 1d**), indicating the high sensitivity of the femtoliter droplet assay. We observed that 100 μM spectinomycin supplementation in the reaction solution enabled us to specifically detect the artificial ribosome translational activity **(Extended Data Fig. 5a and b)**. Then, to evaluate the sensitivity of the assay, we diluted the cell extract containing the artificial ribosomes using the control cell extract and mixed it with the or1-oSD–LacZ reporter and 100 µM spectinomycin. As a result, we successfully detected the translational activity of the artificial ribosomes even at 49 pM (dilution ratio of 10^5^**) (Fig. 2c**). Our Poisson distribution-based calculation suggested that the assay enabled translational activity detection down to the single ribosome level **(Extended Data Fig. 6)**.

### Reconstitution of SSU biogenesis *in vitro*

We tackled again the reconstitution of SSU biogenesis *in vitro* using the two-sided orthogonal translation system and the deep-leaning-assisted automated femtoliter droplet assay. We hypothesized that mimicking *in vivo* ribosome biogenesis would result in *in vitro* ribosome biogenesis. Our experimental scheme was divided into two sequential reactions (**Fig. 3a**). In the first reaction, we aimed at coactivating the transcription of the artificial rRNA operon with or1-oASD and SpcR, the transcription and translation of 21 SSU r-protein genes, and the coordinated assembly in an optimized S150 cell extract, containing the ∼100 accessory factors for ribosome biogenesis and imitating cytoplasmic chemical conditions. The second reaction was designed for detecting nascent artificial SSU translational activity using the or1-oSD–LacZ reporter. We observed no reporter signal during the initial trial for reconstituting SSU biogenesis even using the deep-leaning-assisted automated femtoliter droplet assay, confirming again the difficulty to reconstitute SSU biogenesis **(Extended Data Fig. 7a)**. Then, we thoroughly explored the reaction conditions using a simplex-lattice design and optimized the concentrations of the native ribosomes, the artificial rRNA operon, and 21 SSU r-protein genes. We hypothesized that increasing ribosomal gene concentrations could be beneficial to reconstituting SSU biogenesis as higher DNA concentrations usually produce robust expression profiles^41^ **(Extended Data Fig. 4a)**. However, contrary to our expectations, reducing their concentrations was pivotal and led to slight reporter signal detection (**Fig. 3b**). We conducted a follow-up optimization and successfully optimized the reaction conditions that generated almost saturated reporter signals in the femtoliter droplet assay (**Fig. 3c** **and Extended Data Fig. 7b)**. Using the optimized reaction condition, we detected a strong, reconstituted SSU biogenesis-derived fluorescence signal even in the bulk assay (**Fig. 3d**). This fluorescence signal was stronger than that derived from the nonautonomous iSAT assembly, suggesting that the autonomous *in vitro* ribosome self-replication is highly efficient.

**Fig. 3.**
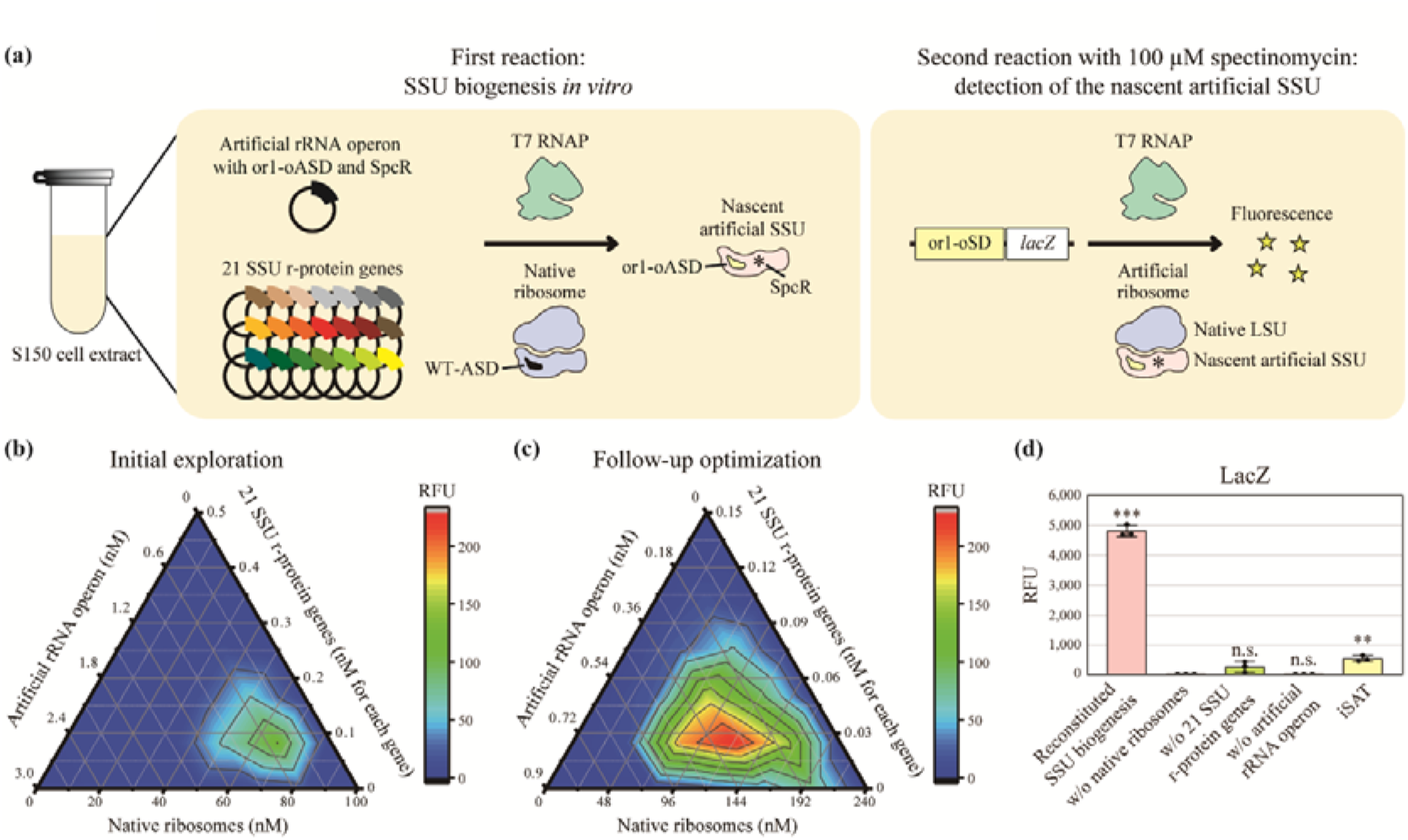
Reconstitution of SSU biogenesis *in vitro*. **(a)** Experimental scheme to reconstitute SSU biogenesis *in vitro*. **(b)** Exploring optimal conditions for the first reaction using a simplex-lattice design. The concentrations of the native ribosomes, the artificial rRNA operon with or1-oASD and C1192U spectinomycin resistance (SpcR), and 21 SSU r-protein genes were 0–100, 0–3, and 0–0.5 nM each, respectively. RFU, mean relative fluorescence unit of the fluorescent droplets. **(c)** Follow-up optimization of the first reaction. The concentrations of the native ribosomes, the artificial rRNA operon, and 21 SSU r-protein genes were 0–240, 0–0.9, and 0–0.15 nM each, respectively. **(d)** Successful detection of the nascent artificial SSU translational activity using the bulk assay under the optimized reaction condition. The concentrations of the native ribosomes, the artificial rRNA operon, and 21 SSU r-protein genes were 80, 0.3, and 0.05 nM each, respectively. RFU, relative fluorescence unit. The data represent the mean ± SD (n = 3). ***, *p* < 0.001; **, *p* < 0.01; n.s., not significant; Dunnett’s test against the negative control without native ribosomes.

### Reconstitution of LSU biogenesis *in vitro*

We moved ahead to reconstitute LSU biogenesis *in vitro*. Our experimental scheme was similar to that for SSU (**Fig. 4a**). The first reaction consisted of coactivating the transcription of an artificial rRNA operon with A2058U clindamycin resistance (CldR)^42^ in the 23S rRNA, the transcription and translation of 33 LSU r-protein genes, and the coordinated assembly in the optimized S150 cell extract. The second reaction was designed for detecting nascent artificial LSU translational activity using the WT-SD–LacZ reporter in the presence of 1.5 mM clindamycin. We expected that detection of the nascent artificial LSU translational activity would be difficult for three reasons: 1) LSU biogenesis is far more complex than SSU biogenesis^2^; 2) the two-sided orthogonal translation system is not available as the nascent artificial LSU requires native SSU for translation, the nascent artificial LSU could thus translate both LacZ and 33 LSU r-proteins; 3) ribosomes with the A2058U CldR mutation retain only ∼30 % of their translational activity in the presence of clindamycin^12^. Surprisingly, a simple exploratory experiment based on the optimized reaction condition for the reconstituted SSU biogenesis enabled us to detect significant fluorescence signals derived from the nascent artificial LSU (**Fig. 4bc**). As expected, the fluorescence signal obtained from the reconstituted LSU biogenesis was lower than that from the reconstituted SSU biogenesis (**Fig. 3d and 4c**). Using an improved LacZ reporter with a modified 5′UTR sequence, we successfully enhanced by 3.8-fold the nascent artificial LSU-derived fluorescence signal (**Fig. 4d**).

**Fig. 4.**
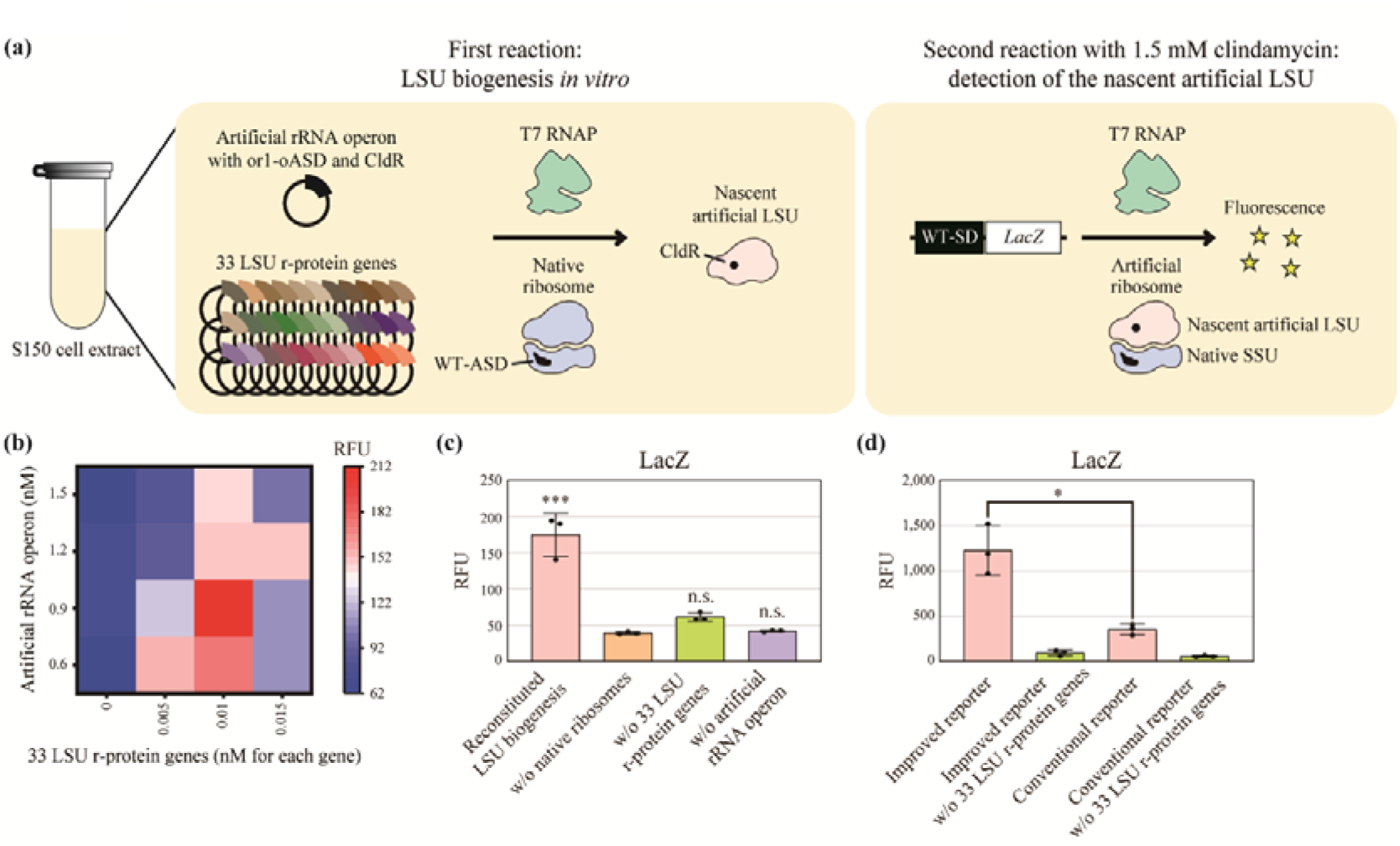
Reconstitution of LSU biogenesis *in vitro*. **(a)** Experimental scheme to reconstitute LSU biogenesis *in vitro*. **(b)** Exploring optimal conditions for the first reaction using the bulk assay. The concentrations of the native ribosomes, artificial rRNA operon with A2058U clindamycin resistance (CldR), and 33 LSU r-protein genes were 80, 0.6–1.5, and 0–0.015 nM each, respectively. RFU, relative fluorescence unit. **(c)** Reproducible detection of the nascent artificial LSU translational activity under the optimized reaction condition. The concentrations of the native ribosomes, the artificial rRNA operon, and 33 LSU r-protein genes were 80, 0.9, and 0.01 nM each, respectively. The data represent the mean ± SD (n = 3). ***, *p* < 0.001; n.s., not significant; Dunnett’s test against the negative control without native ribosomes. **(d)** Improvement of the nascent LSU-derived fluorescence signal using an improved LacZ reporter with a modified 5′UTR sequence. The data represent the mean ± SD (n = 3). *, *p* < 0.05; Welch’s *t*-test.

We tried to obtain further pieces of evidence that ensure the successful reconstitution of SSU and LSU biogenesis. We investigated r-protein production profiles using heavy L-arginine (^13^C_6_, ^15^N_4_) and L-lysine (^13^C_6_, ^15^N_2_) to label nascent r-proteins during the reconstituted SSU and LSU biogenesis. Our mass spectrometric analyses revealed that nascent r-proteins derived from the plasmids encoding r-proteins but not from residual *E. coli* chromosomal fragments or mRNAs in the S150 cell extracts **(Extended Data Fig. 8a)**. Furthermore, we conducted a reconstitution experiment using a mutant r-protein gene encoding uS12 K43T, which confers streptomycin resistance (StrR) to ribosomes^28, 43^. We observed that the reporter signals remained unaffected by streptomycin only when we used the mutant r-protein gene as a starting material **(Extended Data Fig. 8b)**. In addition, a previous study showed that most r-proteins do not exchange between ribosomes^44^. Taken together, we concluded that the nascent artificial ribosomes consisted of nascent rRNAs and r-proteins.

Finally, we investigated whether LSU and SSU could be reconstituted in a single reaction. We coactivated the transcription of an artificial rRNA operon with or1-oASD, SpcR in the 16S rRNA, and CldR in the 23S rRNA, and the production of 54 r-proteins in the optimized S150 cell extract **(Extended Data Fig. 9a)**, leading to the successful reconstitution of both LSU and SSU biogenesis in a single reaction space **(Extended Data Fig. 9b)**. Therefore, we finally succeeded in reconstituting the entire ribosome biogenesis process *in vitro*.

### Conclusions

In this study, we achieved the reconstitution of the entire ribosome self-replication process outside a living cell, which has been one of the largest long-standing challenges in synthetic biology. The reconstituted *in vitro* ribosome biogenesis would provide us with more freedom in controlling the process of ribosome biogenesis. Therefore, this achievement would pave the way to reveal fundamental principles underlying ribosome biogenesis and to elucidate the mechanisms of ribosomopathies^24^. Furthermore, our achievement brings the bottom-up creation of self-replicating artificial cells within reach as life scientists now have successfully activated *in vitro* all the processes necessary for the autonomous central dogma, i.e., DNA replication^45, 46^, transcription^47^, translation^48, 49^, and in this study, ribosome biogenesis. Finally, the nature of our platform (autonomous ribosome assembly *in vitro* from DNA and the lack of cell viability constraints) opens new opportunities for the high-throughput and unconstrained creation of artificial ribosomes with altered or enhanced properties^8–12^, significantly expanding types of polymers available to humankind.

## Extended data figures/tables

**Extended Data Fig. 1.**
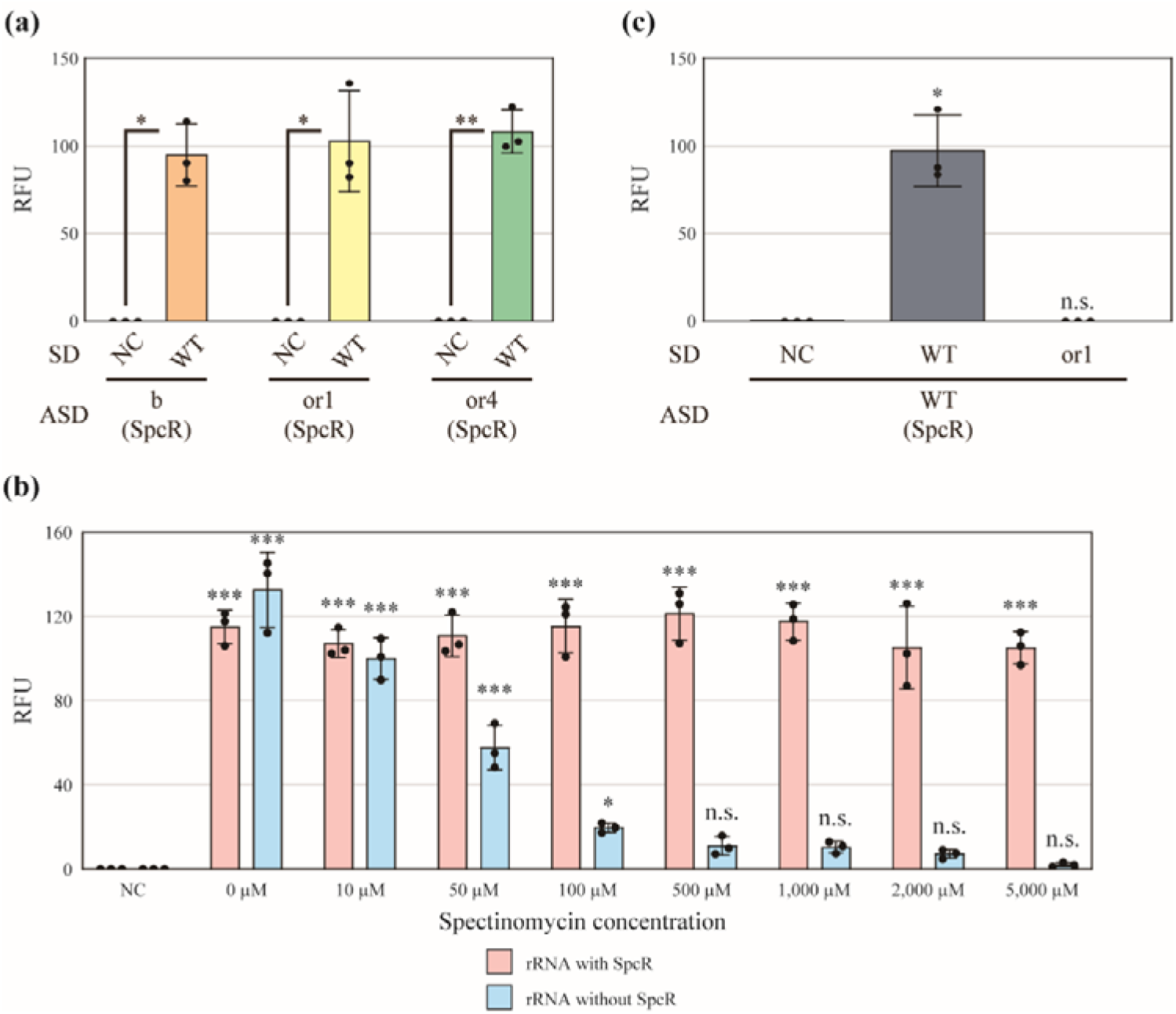
Supporting experiments for screening oSD·oASD pairs with two-sided orthogonality. **(a)** Functional cell extract preparation. S12 Cell extracts were prepared using *E. coli* expressing an artificial rRNA operon with oASD (b, or1, or or4) and C1192U spectinomycin resistance (SpcR). The cell extracts were mixed with a WT-SD–LacZ reporter. The cell extracts that generated strong fluorescence signals were considered functional. RFU, relative fluorescence unit; NC, negative control without the reporter. The data represent the mean ± SD (n = 3). *, *p* < 0.05; **, *p* < 0.01; Welch’s *t*-test. **(b)** Optimizing spectinomycin concentrations. S12 cell extracts were prepared using BL21 Star^TM^ (DE3) *lacZ::frt* expressing an rRNA operon with WT-ASD and/or SpcR. The cell extracts were mixed with the WT-SD–LacZ reporter at various spectinomycin concentrations. We concluded that 5 mM spectinomycin completely inactivated native ribosomes but left the artificial ribosomes with SpcR unaffected. We used spectinomycin at 5 mM in the following experiments if not specified. NC, negative control without the reporter. The data represent the mean ± SD (n = 3). *, *p* < 0.05; ***, *p* < 0.001; n.s., not significant; Dunnett’s test against NC. **(c)** A follow-up control experiment to verify two-sided orthogonality of the or1-oSD·oASD pair. A cell extract was prepared using *E. coli* expressing an artificial rRNA operon with WT-ASD and SpcR. The extract was mixed with the WT-SD–LacZ or the or1-oSD–LacZ reporter in the presence of spectinomycin. The cell extract generated no fluorescence signal when mixed with the or1-oSD–LacZ reporter, indicating that SpcR did not contribute to the fluorescence signal observed in Fig. 1g. NC, negative control without a reporter. The data represent the mean ± SD (n = 3). *, *p* < 0.05; n.s., not significant; Dunnett’s test against NC.

**Extended Data Fig. 2.**
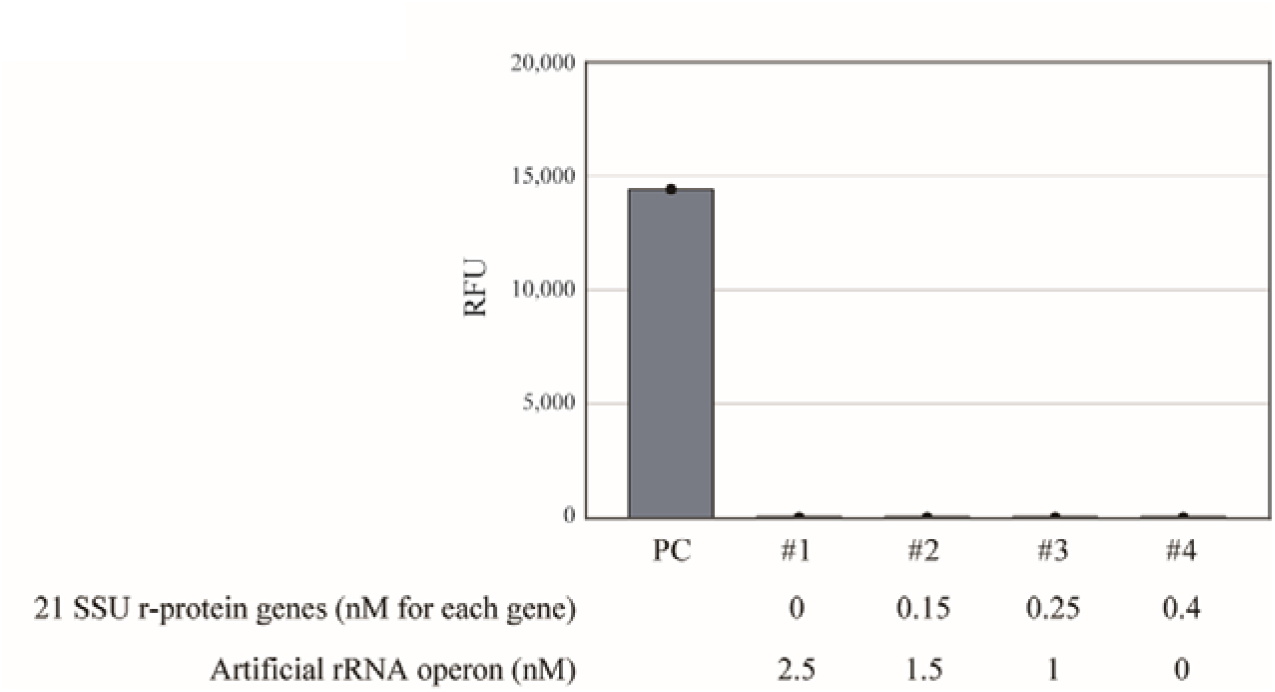
An initial trial to reconstitute SSU biogenesis. Experimental scheme to reconstitute SSU biogenesis *in vitr*o is shown in Fig. 3a. Briefly, the first reaction is designed for coactivating the transcription of the artificial rRNA operon with or1-oASD and SpcR, the transcription and translation of 21 SSU r-protein genes, and coordinated assembly in an optimized S150 cell extract, containing the ∼100 accessory factors for ribosome biogenesis and imitating cytoplasmic chemical conditions. The second reaction aimed at detecting the translational activity of nascent artificial SSU using the or1-oSD–LacZ reporter. Exploring several reaction conditions did not generate any fluorescence signals derived from nascent SSU. The concentrations of the native ribosomes, the artificial rRNA operon with or1-oASD and C1192U spectinomycin resistance (SpcR), and 21 SSU r-protein genes were 20, 0–2.5, and 0–0.4 nM each, respectively. RFU, relative fluorescence unit; PC, positive control using native ribosomes and the WT-SD–LacZ reporter.

**Extended Data Fig. 3.**
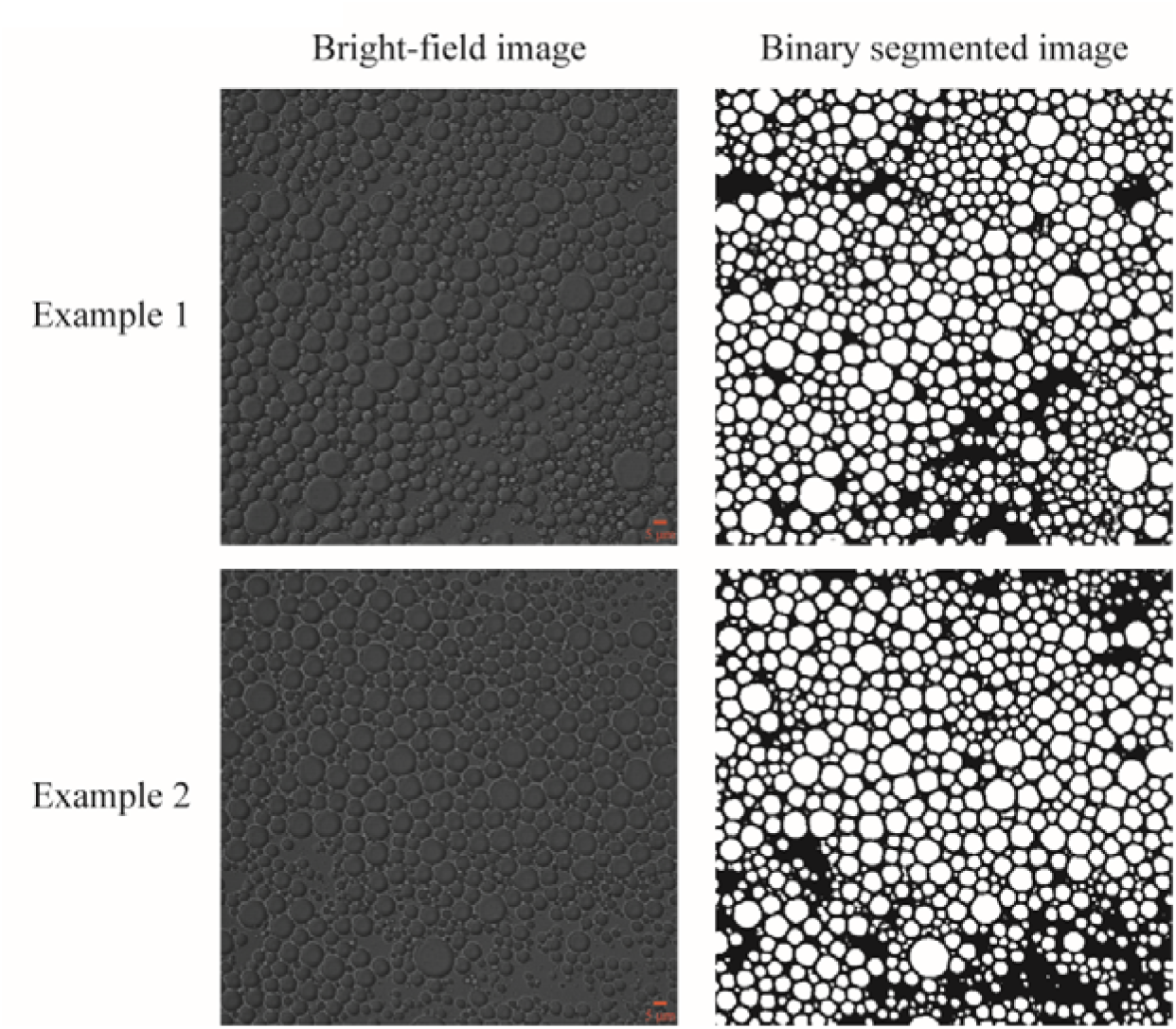
Binary segmentation by the trained U-Net deep-learning model. Bright-field images of the droplets (left panels) were processed into binary segmented images (right panels; droplet or background) with >90 % accuracy (number of correctly classified pixels/total number of pixels) using the trained U-Net deep-learning model. In the right panels, white and black regions indicate the droplets and background, respectively. Scale bars = 5 μm.

**Extended Data Fig. 4.**
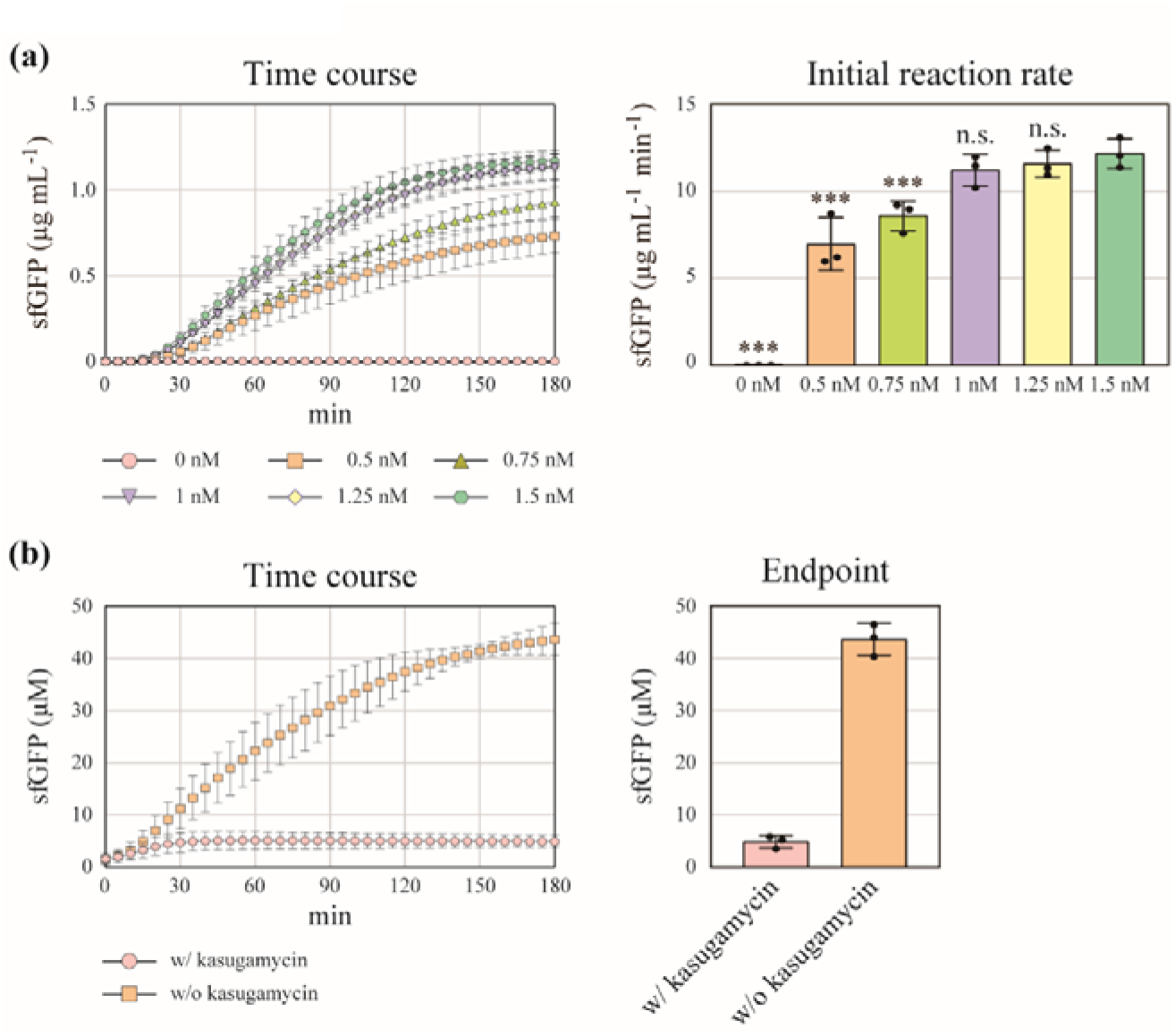
Estimation of the artificial ribosome concentration with or1-oASD and C1192U spectinomycin resistance (SpcR). **(a)** Determining non-rate-limiting concentrations of the or1-oSD–sfGFP reporter. The S12 cell extract was prepared using BL21 Star^TM^ (DE3) *lacZ::frt* expressing the artificial rRNA operon with or1-oASD and SpcR. The cell extract was mixed with spectinomycin and 0–1.5 nM of the or1-SD–sfGFP reporter. We found that transcription was not rate-limiting when reporter concentrations were over 1 nM. The data represent the mean ± SD (n = 3). ***, *p* < 0.001; n.s., not significant; Dunnett’s test against 1.5 nM. **(b)** Quantification of the number of the artificial ribosomes with or1-oASD and SpcR. The cell extract was mixed with spectinomycin and 1 nM of the or1-oSD–sfGFP reporter. In the presence of kasugamycin, the concentration of produced sfGFP equals that of the artificial ribosomes. The estimated concentration of the artificial ribosomes was 4.9 µM. The data represent the mean ± SD (n = 3).

**Extended Data Fig. 5.**
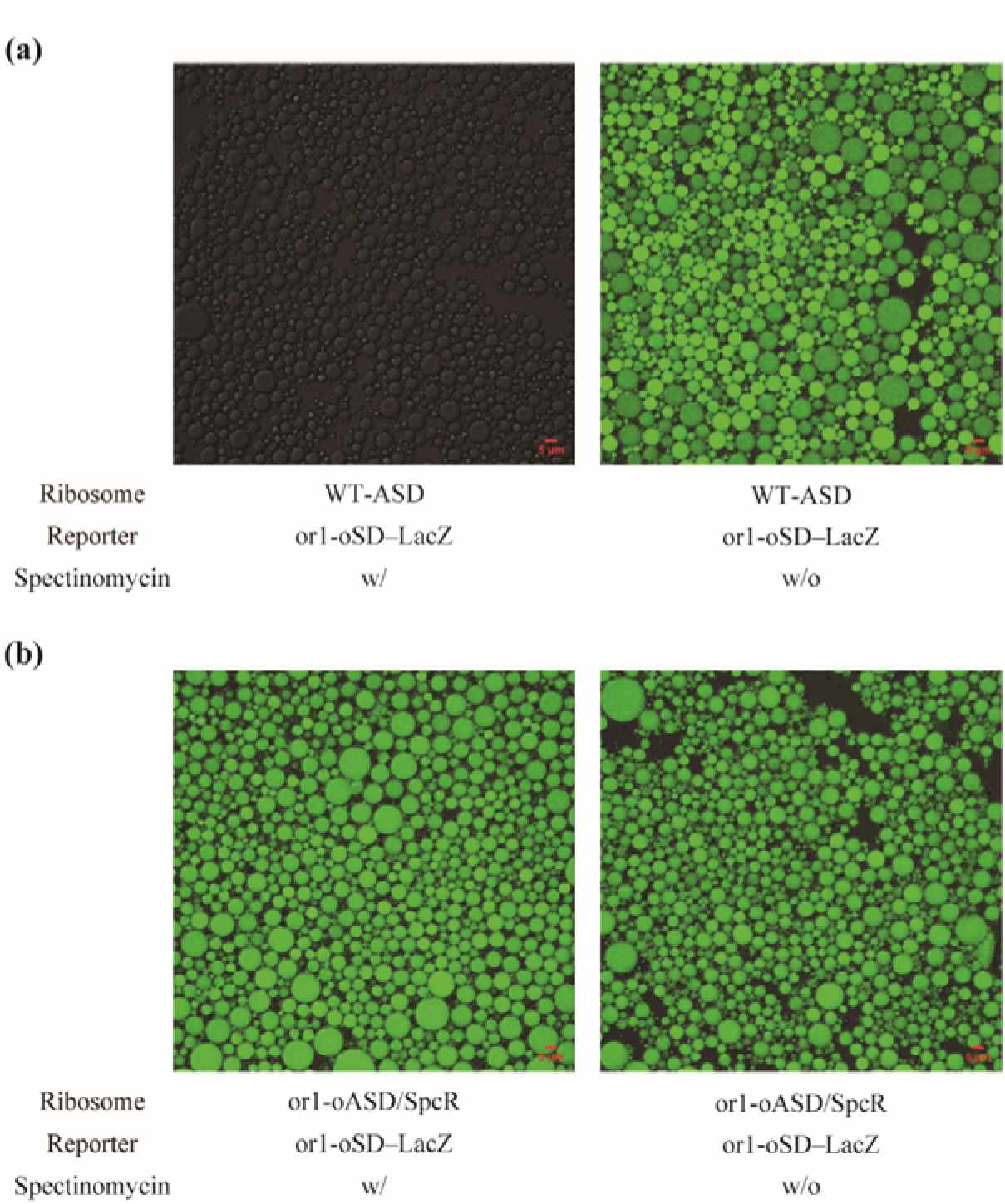
Native ribosome deactivation by spectinomycin in the femtoliter droplet assay. **(a)** Representative micrographs of droplets after the femtoliter droplet assay using control S12 cell extracts containing only native ribosomes. The concentration of the or1-oSD–LacZ reporter was 5 nM. The droplets emitted strong fluorescence without spectinomycin as the femtoliter droplet assay is so sensitive that a very weak interaction between the native ribosomes and the or1-oSD–LacZ reporter could be detected. The addition of 100 μM spectinomycin eliminated this nonspecific fluorescence signal. Scale bars = 5 μm. **(b)** Representative micrographs of droplets after the femtoliter droplet assay using S12 cell extracts containing artificial ribosomes with or1-oASD and C1192U spectinomycin resistance (SpcR). The concentration of the or1-oSD–LacZ reporter was 5 nM. The droplets emitted strong fluorescence with or without 100 μM spectinomycin. Scale bars = 5 μm.

**Extended Data Fig. 6.**
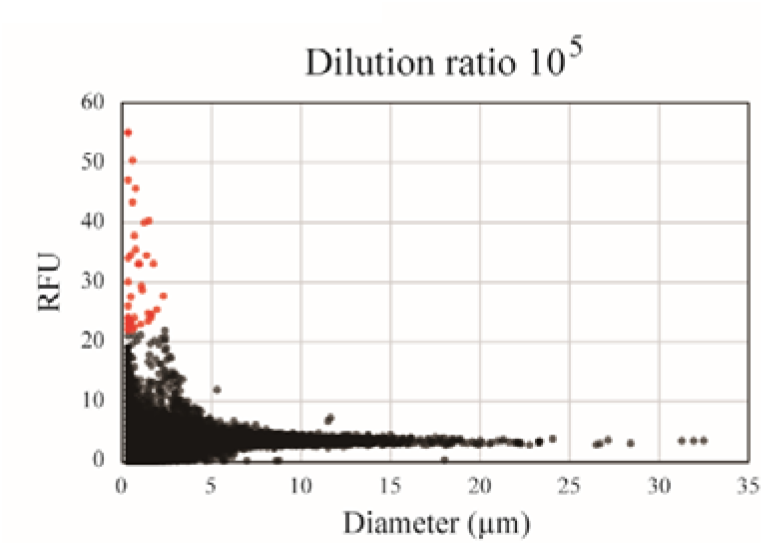
Single-ribosome-level detection of the artificial ribosome translational activity. Scatter plot of the mean relative fluorescence unit (RFU) against the diameter of each droplet was generated using the dataset of Fig. 2c. Droplets over the threshold (the mean RFU ≥ 22) are shown in red. From the Poisson distribution formula, most of the fluorescent droplets (89 %) were estimated to contain only one artificial ribosome.

**Extended Data Fig. 7.**
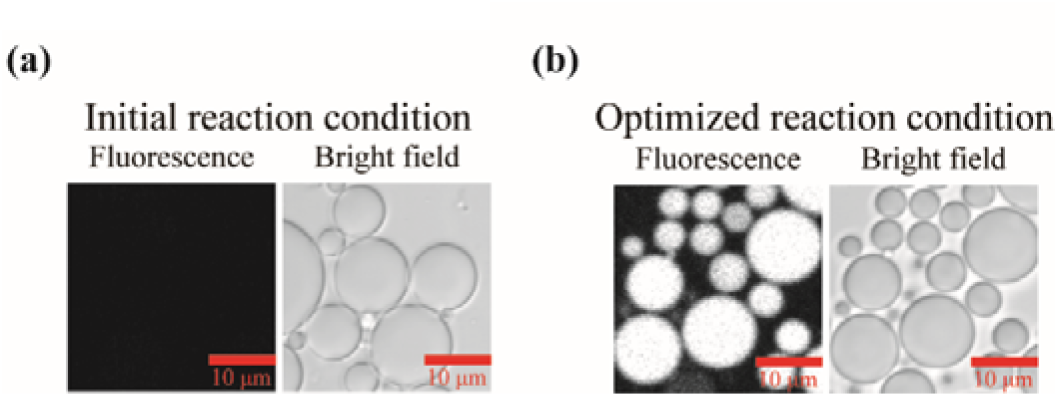
Representative images of the femtoliter droplet assay. **(a)** A representative micrograph of droplets in the initial trial. In the first reaction, the concentrations of the native ribosomes, the artificial rRNA operon with or1-oASD and C1192U spectinomycin resistance (SpcR), and 21 SSU r-protein genes were 20, 1, and 0.25 nM each, respectively. Scale bars = 10 μm. **(b)** A representative micrograph of droplets in the optimized reaction condition. The concentrations of the native ribosomes, the artificial rRNA operon, and 21 SSU r-protein genes were 80, 0.3, and 0.05 nM each, respectively. Scale bars = 10 μm.

**Extended Data Fig. 8.**
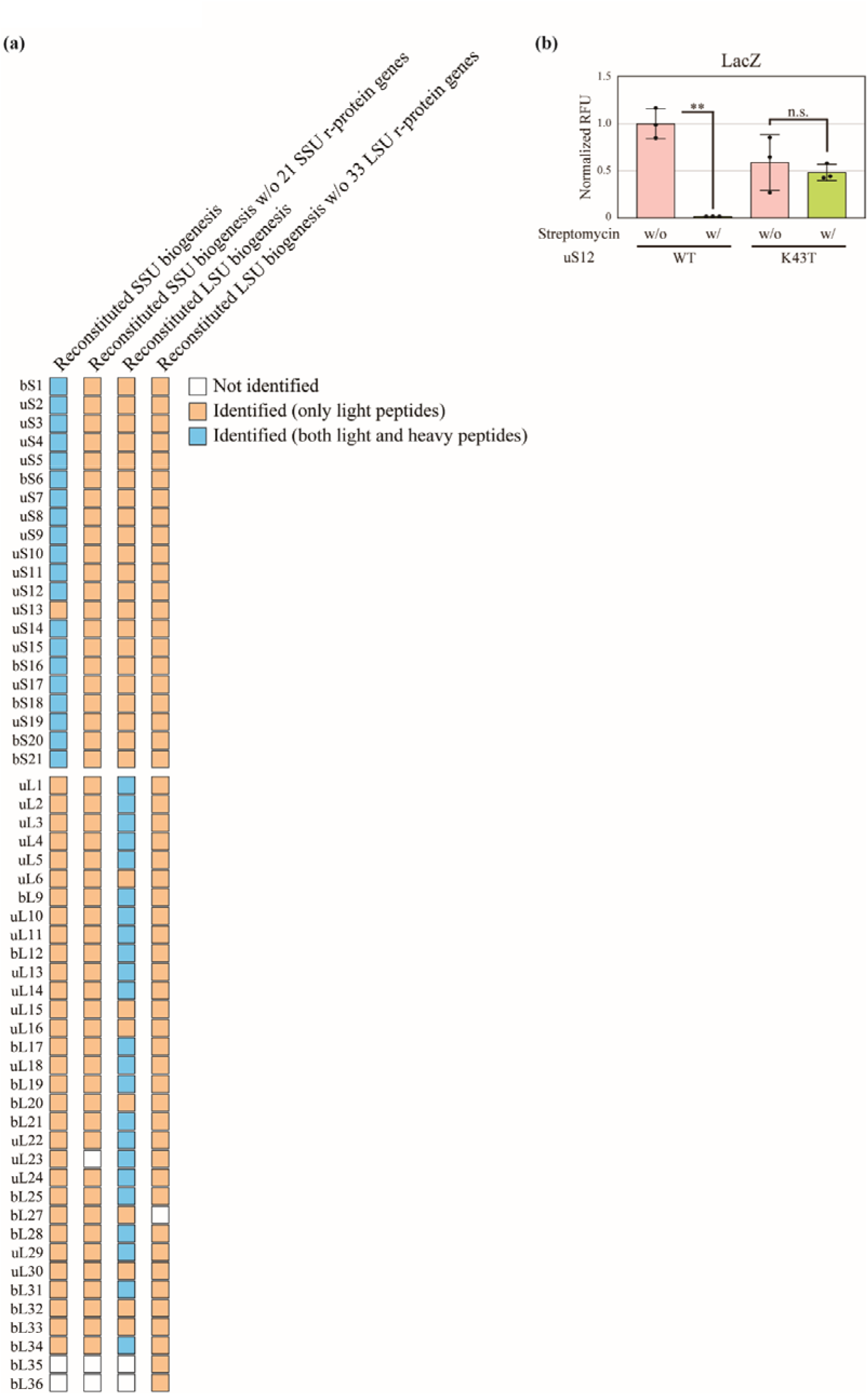
Supporting pieces of evidence for the *in vitro* SSU and LSU biogenesis. **(a)** Detection of pre-existing and nascent r-proteins. We replaced unlabeled (light) L-arginine and L-lysine with stable isotope-labeled (heavy) L-arginine (^13^C_6_, ^15^N_4_) and L-lysine (^13^C_6_, ^15^N_2_) to label nascent r-proteins in the reaction solutions during the reconstituted SSU or LSU biogenesis (Fig. 3 and 4). As negative controls, we omitted the r-protein genes from the reaction solutions. Certain r-proteins were not identified (e.g., bL35 and bL36 in the reconstituted SSU biogenesis) because r-proteins are very small and difficult targets for proteomics. The heavy peptides of certain r-proteins were not identified (e.g., uS13 in the reconstituted SSU biogenesis); the absence of heavy peptides does not mean the absence of nascent proteins because of the stochastic nature of the protein identification algorithm. **(b)** Direct evidence for the incorporation of newly synthesized r-proteins into nascent ribosomes. The concentrations of the native ribosomes, the artificial rRNA operon with or1-oASD and C1192U spectinomycin resistance (SpcR), and 21 SSU r-protein genes were 80, 0.3, and 0.05 nM each, respectively. A mutant r-protein gene encoding uS12 K43T was used instead of an r-protein gene encoding native uS12. The uS12 K43T mutation confers streptomycin resistance to SSU. The translational activity of the nascent artificial SSU was detected using the or1-oSD–LacZ reporter in the presence of spectinomycin and streptomycin. NC, negative control without 21 SSU r-protein genes. The data represent the normalized relative fluorescence unit (RFU) in the bulk assay and are shown as the mean ± SD (n = 3). **, *p* < 0.01; Welch’s *t*-test.

**Extended Data Fig. 9.**
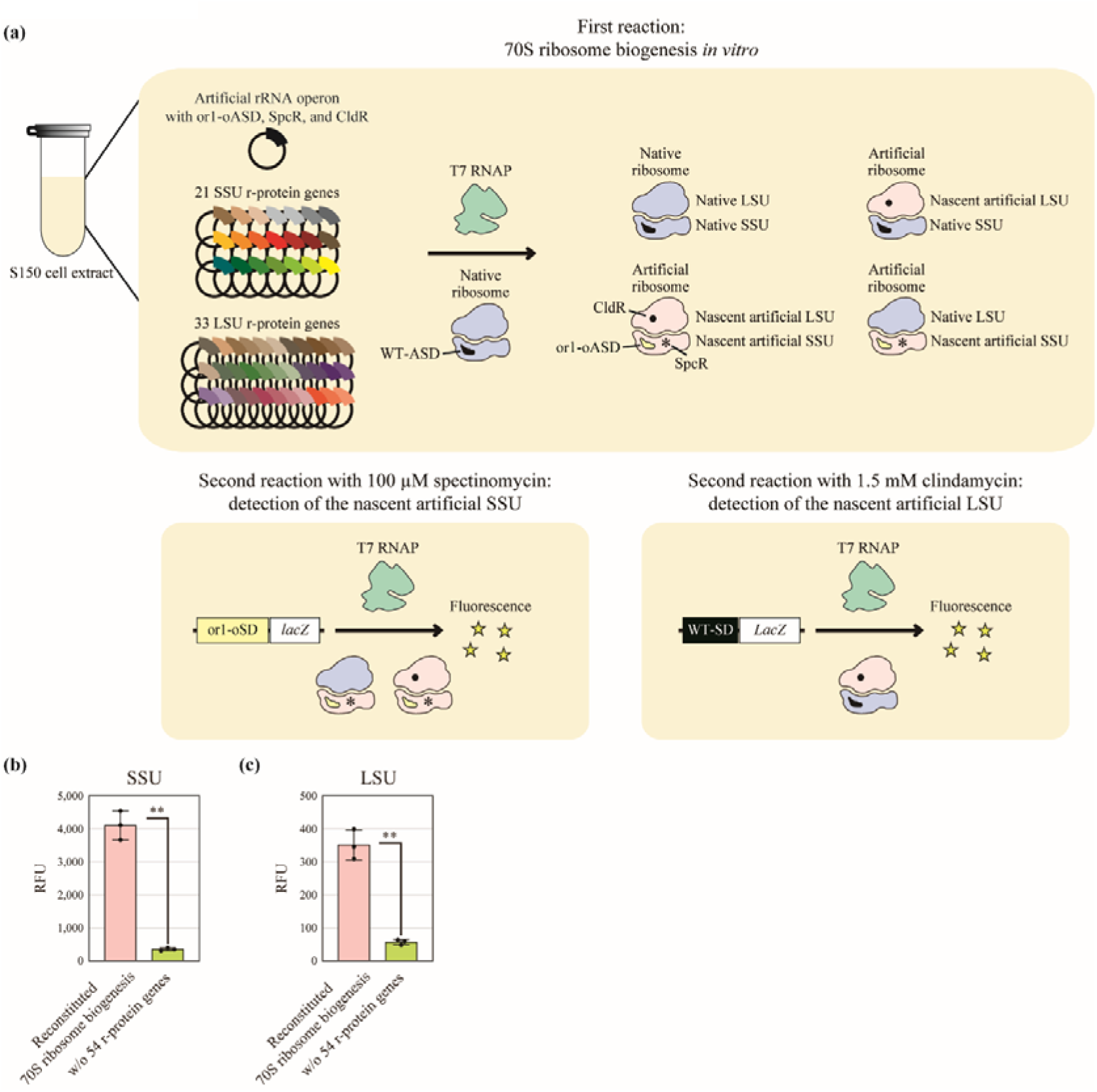
*In vitro* reconstitution of the entire ribosome biogenesis process in a single reaction. **(a)** Experimental scheme to reconstitute both SSU and LSU biogenesis *in vitro* in a single reaction. **(b and c)** Successful detection of the nascent artificial SSU and LSU translational activity using the bulk assay under the optimized reaction condition. The concentrations of the native ribosomes, artificial rRNA operon with or1-oASD, C1192U spectinomycin resistance (SpcR), and A2058U clindamycin resistance (CldR), and 54 r-protein genes were 80, 0.9, and 0.01 nM each, respectively. The data represent the mean ± SD (n = 3). **, *p* < 0.01; Welch’s *t*-test.

## Methods

### Strains and plasmids

The chromosomal *lacZ* gene of the BL21 Star^TM^ (DE3) (Thermo Fisher Scientific, Waltham, MA, USA) was disrupted by Red-mediated recombination^50^. Briefly, pKD46 encoding phage λ-Red recombinase was transformed into the *E. coli* cells. The transformants were grown in 50 mL of SOC medium with 100 μg/ml ampicillin (Viccillin^®^ for injection, Meiji Seika Pharma, Tokyo, Japan) and 10 mM L-(+)-arabinose (Nacalai Tesque, Kyoto, Japan). The fragment of the kanamycin-resistance gene (*kmr*) was amplified using primers (H1P1 forward primer, 5′-GAAATTGTGAGCGGATAACAATTTCACACAGGAAACAGCTGTGTAGGCTGGAGCTGCTTC-3′, and P4H2 reverse primer, 5′-TTACGCGAAATACGGGCAGACATGGCCTGCCCGGTTATTAATTCCGGGGATCCGTCGACC-3′) from pKD13, and introduced into the *E. coli* cells by electroporation. The electroporated cells were grown on an LB agar plate with 50 μg/ml of kanamycin monosulfate (Nacalai Tesque) to select Km^R^ transformants. The resulting strain is described as BL21 Star^TM^ (DE3) *lacZ::kmr*. The FLP helper plasmid, pCP20, was transformed into the BL21 Star^TM^ (DE3) *lacZ::kmr* to eliminate the *kmr* gene. As pCP20 harbors a temperature-sensitive replicon and shows thermal induction of FLP synthesis, the transformants were cultured nonselectively at 37 °C and tested for the loss of antibiotic resistance. The resulting strain is described as BL21 Star^TM^ (DE3) *lacZ::frt*.

The *rrnB* rRNA operon was inserted into pET-41a(+). Genes encoding 54 r-proteins were cloned from the *E. coli* DH5α genome. In this study, we included bS1 in r-proteins, which works more as a translational factor than a structural component, because additional bS1 could improve protein yields^51–54^. The expression of the genes encoding 54 r-proteins was regulated by the *pT7CONS*^55^ and *EpsA20*^56^ sequences, improving transcription and translation efficiencies. *pT7CONS* and *EpsA20* were also used to construct an improved LacZ reporter. Mutations in genes encoding rRNAs and r-proteins were introduced by mutagenic primer-based PCR.

The strains and plasmids used in this study are listed in **Supplementary Information 2**.

### Sonicated S12 cell extract preparation

Sonicated S12 cell extracts were prepared as previously described with some modifications^57^. Briefly, *E. coli* cells were grown in 200 mL of 2 × YPTG medium at 37 °C and pelleted by centrifugation. The cell pellets were resuspended in buffer A. The suspended cells were disrupted by a Q125 Sonicator (Qsonica, Newtown, CT, USA) at a frequency input of 20 kHz and amplitude of 50 %. The sonication energy input was 500 J for 1 mL cell suspension. The cell extract was centrifuged at 4 °C and 12,000 *g* for 10 min, and the supernatant was collected. The obtained cell extract was flash-frozen in liquid nitrogen and preserved at −80 °C until further use.

### French press cell extract preparation

French press S30 cell extracts were prepared based on previous reports with some modifications^4, 5, 58^. Briefly, *E. coli* cells were grown in 1 L of 2 × YPTG medium at 37 °C and pelleted by centrifugation. The cell pellets were resuspended in buffer A (20 mM Tris-HCl, 100 mM NH_4_Cl, 10 mM MgCl_2_, 0.5 mM EDTA, and 2 mM DTT, pH = 7.2). Halt Protease Inhibitor Cocktail (Thermo Fisher Scientific) and RNase Inhibitor (QIAGEN) were added to the suspension. The cells were disrupted using an EmulsiFlex-C5 homogenizer (Avestin, Ottawa, Canada) with a single pass at a pressure of 20,000 psi. RNase Inhibitor and DTT were added to the cell extracts followed by centrifugation at 4 °C, 30,000 *g* for 30 min twice. The collected supernatant was dialyzed four times against the iSAT buffer (50 mM HEPES-KOH, 10 mM magnesium glutamate, 200 mM potassium glutamate, 2 mM DTT, 1 mM spermidine, and 1 mM putrescine), imitating cytoplasmic chemical conditions^58^. For clarification and concentration, the cell extract was centrifuged at 4,000 *g* for 10 min in a Centriprep^®^ 3K device (EMD Millipore, Burlington, MA, USA). The obtained cell extract was flash-frozen in liquid nitrogen and preserved at −80 °C until further use.

French press S150 cell extracts were prepared as previously described with some modifications^4, 5^. Briefly, BL21 Star™ (DE3) *lacZ::frt* harboring pT7_WT-ASD_rRNA was grown in 1 L of 2 × YPTG medium with 50 µg/mL of kanamycin at 37 °C until the OD_600_ reached 0.5. The cells were incubated with 0.1 mM isopropyl-β-D-thiogalactopyranoside (IPTG, Nacalai Tesque). Then, the cells were disrupted using an EmulsiFlex-C5 homogenizer (Avestin) with a single pass at a pressure of 20,000 psi. The cell extracts were centrifuged at 30,000 *g* for 30 min at 4 °C. The collected supernatants were centrifuged at 90,000 *g* for 21 h at 4 °C. The collected supernatants were further centrifuged at 150,000 *g* for 3 h at 4 °C. Then, the collected supernatants were dialyzed using the iSAT buffer. The cell extracts were concentrated using Amicon Ultra-15 3 kDa cutoff (Merck Millipore, Burlington, MA, USA). The obtained cell extract was flash-frozen in liquid nitrogen and preserved at −80 °C until further use.

### Cell extract preparation containing ribosomes with artificial rRNAs

A plasmid encoding an artificial rRNA operon was introduced into the BL21 Star^TM^ (DE3) *lacZ::frt*. The transformant was grown in a 2 × YPTG medium with 50 µg/mL of kanamycin at 37 °C until the OD_600_ reached 0.7. The cultured cells were incubated with 0.1 mM IPTG (Nacalai Tesque) for 3 h. The cell extracts were prepared as described above.

### Cell-free transcription and translation (CF-TXTL)

CF-TXTL was performed according to a previous report with modifications^4^. *E. coli* ribosomes were purchased from New England BioLabs (Ipswich, MA, USA). T7 RNA polymerase (T7 RNAP, New England BioLabs) was added to a final concentration of 0.8 U/μL. T7 RNAP was not added when we used cell extracts derived from IPTG-induced BL21 Star^TM^ (DE3) or its derivative strains. The reporter plasmid concentration was 1.5 nM. The sfGFP or LacZ reporter expression was induced by IPTG at a final concentration of 2 mM. We used 5-chloromethylfluoresecein di-β-D-galactopyranoside (CMFDG; Invitrogen, Waltham, MA, USA) as a substrate of LacZ at a final concentration of 33 μM. CF-TXTL was conducted using 15 μL reaction solutions at 37 °C in a 96-well plate (polystyrene, solid bottom, half area, black-walled, Greiner Bio-One International GmbH, Kremsmünster, Austria). The reporter signals were quantified using fluorescence microplate readers, Fluoroskan Ascent FL^TM^ (Thermo Fisher Scientific) or Infinite^®^ 200 PRO (TECAN, Männedorf, Switzerland), at λ_ex_ = 485 nm and λ_em_ = 535 nm. For native ribosome deactivation, spectinomycin (FUJIFILM Wako Pure Chemical Corporation, Osaka, Japan), streptomycin (FUJIFILM Wako Pure Chemical Corporation), or clindamycin (Abcam, Cambridge, UK) were used at final concentrations of 5 mM, 10 μg/mL, or 1.5 mM, respectively. For the ribosome concentration quantification, kasugamycin (FUJIFILM Wako Pure Chemical Corporation) was added to 2 mM 15 min after the beginning of CF-TXTL as previously reported^59^. The fluorescence intensity was kinetically measured after adding kasugamycin, and the background fluorescence intensity was subtracted. Kasugamycin is an antibiotic originally isolated from *Streptomyces kasugaensis* that blocks translation initiation by preventing the ribosomal subunit association. However, it did not affect translating or stalled 70S ribosomes^60, 61^. The constituents of the CF-TXTL reaction solutions used in this study are summarized in **Supplementary Information 3**.

The *in vitro* reconstitution of SSU biogenesis was performed as follows. In the first reaction, the CF-TXTL solutions based on the S150 cell extracts were mixed with the native ribosomes, the artificial rRNA operon with or1-oASD and C1192U SpcR, and 21 SSU r-protein genes. We used S150 cell extracts to enable native ribosome concentration control. The solutions were incubated at 37 °C for 180 min. The reaction conditions were optimized using a simple lattice design **(Supplementary Information 4)**. In the second reaction, the resulting CF-TXTL solutions were mixed with pT7_or1-oSD_LacZ, CMFDG, 100 µM spectinomycin, and an additional 15 µL of the CF-TXTL solutions based on the S150 cell extracts. The fluorescence of the reaction solutions was measured by a bulk assay or a femtoliter droplet assay. In the bulk assay, the reporter signals were kinetically measured at 37 °C using Infinite^®^ 200 PRO (TECAN) at λ_ex_ = 485 nm and λ_em_ = 535 nm. The femtoliter droplet assay was carried out as described below.

The *in vitro* reconstitution of LSU biogenesis was performed as follows. In the first reaction, the CF-TXTL solutions based on the S150 cell extracts were mixed with the native ribosomes, the artificial rRNA operon with or1-oASD, SpcR, and A2058U CldR, and 33 LSU r-protein genes. The solutions were incubated at 37 °C for 180 min. The reaction conditions were optimized by varying the concentrations of the artificial rRNA operon and 33 LSU r-protein genes **(Supplementary Information 4)**. In the second reaction, the resulting CF-TXTL solutions were mixed with pT7_WT-SD_LacZ or pT7PCONS_EpsA20_WT-SD_lacZ, CMFDG, clindamycin, and an additional 15 µL of the CF-TXTL solutions based on the S150 cell extracts. The fluorescence of the reaction solutions was kinetically measured at 37 °C using Infinite^®^ 200 PRO (TECAN) at λ_ex_ = 485 nm and λ_em_ = 535 nm.

The *in vitro* reconstitution of the entire ribosome biogenesis process was conducted according to the protocol described above with minor modifications. In the first reaction, the concentrations of the native ribosomes, the artificial rRNA operon with or1-oASD, SpcR, and CldR, and 54 r-protein genes were 80, 0.9, and 0.01 nM each, respectively. In the second reaction, pT7_or1-oSD_LacZ and spectinomycin were used for the detection of the nascent artificial SSU, and pT7PCONS_EpsA20_WT-SD_lacZ and clindamycin were used for the detection of the nascent artificial LSU. The reaction solution fluorescence was kinetically measured at 37 °C using Infinite^®^ 200 PRO (TECAN) at λ_ex_ = 485 nm and λ_em_ = 535 nm.

The iSAT assembly was performed according to previous reports^4, 5, 58^ with some modifications. Briefly, in the first reaction, the CF-TXTL solutions based on the S150 cell extracts were mixed with 100 nM total protein of 70S ribosome (TP70) and 0.3 nM of the artificial rRNA operon with or1-oASD and SpcR. The solutions were incubated at 37 °C for 180 min. In the second reaction, the resulting CF-TXTL solutions were mixed with pT7_or1-oSD_LacZ, CMFDG, spectinomycin, and an additional 15 µL of the CF-TXTL solutions based on the S150 cell extracts. The reaction solution fluorescence was kinetically measured at 37 °C using Infinite^®^ 200 PRO (TECAN) at λ_ex_ = 485 nm and λ_em_ = 535 nm.

The parameter values described above were tuned experiment-dependently and are specified in the figure legends.

### Femtoliter droplet assay

An oil mixture was composed of light mineral oil (Sigma-Aldrich Corporation, St. Louis, MO, USA), 4.5 % sorbitan monooleate (Nacalai Tesque), and 0.5 % Triton^®^ X-100 (Nacalai Tesque), as previously described^62, 63^. The CF-TXTL reaction solutions were mixed with the oil mixture and tapped twenty times in microtubes (Maruemu Corporations, Osaka, Japan). The emulsions were incubated at 37 °C. The bright-field and fluorescence images of droplets were obtained using a confocal fluorescence microscope LSM700 (Carl Zeiss AG, Oberkochen, Germany). The 488 nm laser was focused using an oil immersion objective (Plan-Apochromat 40×/1.4 Oil DIC M27, Carl Zeiss AG) with immersion oil (Immersol^TM^ 518F, Carl Zeiss AG).

### Deep-learning-assisted automated femtoliter droplet assay

It is a difficult task to extract features from a large number of droplets; hence, we devised a deep-leaning-assisted automated analysis pipeline for a scalable and objective femtoliter droplet assay. We aimed to develop an analysis pipeline enabling area and centroid extraction of each droplet from the bright-field images and the fluorescence intensity of each droplet from the corresponding fluorescence images. In the beginning, we produced positive control fluorescent droplets using purified LacZ (FUJIFILM Wako Pure Chemical Corporation) and CMFDG and generated 15 sets of bright-field and corresponding fluorescence images containing 27580 fluorescent droplets in total. We trained an ilastik^64^ pixel classification model and processed the positive control fluorescence images into binary segmented images (droplet or background) as ground truth. We used a convolutional neural network architecture called U-Net^39^ to build a binary segmentation model. We used the FastAI library^65^ under an Anaconda virtual environment (Python 3.7, torch==1.4.0+cpu, torchvision==0.5.0+cpu). The model was trained using 13 sets of ground-truth binary segmented images and the corresponding bright-field images, and the remaining two sets of images were used as test data. We specified an encoder network, Resnet34, and a weight-decay of 1e-2. We searched for a fitting learning rate using the learn.lr_find() method, and picked a learning rate of 1e-4. The model was trained using the fit_one_cycle() method for 20 epochs at slice(1e-4) and pct_start=0.3. We unfroze all layers and searched for a learning rate again. The whole model was trained using the fit_one_cycle() method for 100 epochs at slice(1e-4) and pct_start=0.3. As a result, the accuracy (number of correctly classified pixels/total number of pixels) reached >90 % using the test data **(Extended Data Fig. 3)**. The trained U-Net deep-learning model was used to process bright-field droplet images into binary segmented images, in which white and black regions indicate the droplets and background, respectively. The binary segmented images were provided for particle analysis using ImageJ, and the particle analysis results were redirected to corresponding fluorescence images. Using the deep-learning-assisted automated analysis pipeline, we could automatically obtain the area, mean fluorescence intensity, minimum fluorescence intensity, maximum fluorescence intensity, integrated density, and centroid of each droplet in a scalable and objective manner. The codes were described in **Supplementary Information 5**.

### Sensitivity calculation of the deep-learning-assisted automated femtoliter droplet assay

We roughly estimated the sensitivity of our femtoliter droplet assay using the data at 49 pM artificial ribosomes (dilution ratio 10^5^) (**Fig. 2c** and **Extended Data Fig. 6)**. In this experiment, a droplet with a diameter of 1 µm was expected to contain an average of 1.54 × 10^−2^ ribosomes. From the Poisson distribution formula, the probability that a 1-µm droplet would contain k ribosomes was expressed as follows:

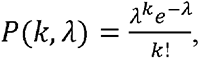

where k is the number of ribosomes and λ is 1.54 × 10^−2^. According to this formula, the ratios of droplets that contain zero, one, or two or more ribosomes were 0.9847, 0.0152, or 0.0001, respectively.

In the data at 49 pM artificial ribosomes (dilution ratio 10^5^), we observed 15544 droplets with a diameter of 0.5–1.5 μm, and the number of fluorescent droplets among them was 17. The observed ratio of the fluorescent droplets was 0.0011. Taken together, most of the fluorescent droplets (89%) were estimated to contain only a single artificial ribosome.

### Mass spectrometric analysis

The proteomic analysis was carried out as previously described with modifications^66^. Briefly, the CF-TXTL reaction solutions were reduced by 50 mM dithiothreitol and modified with 50 mM iodoacetamide. For stable isotope labeling^67^, we used CF-TXTL reaction solutions with 20 amino acid mixtures containing stable isotope-labeled (heavy) L-arginine (^13^C_6_, ^15^N_4_) and L-lysine (^13^C_6_, ^15^N_2_) (Thermo Fisher Scientific) instead of unlabeled (light) L-arginine and L-lysine. The proteins were digested with sequencing-grade modified trypsin (Promega Corporation, Madison, WI, USA). The peptides were analyzed using a nano LC–MS system (UltiMate^TM^ 3000 RSLCnano and Orbitrap Exploris^TM^ 240) equipped with an Aurora UHPLC column (AUR2-25075C18A; IonOpticks, Fitzroy, Australia). A gradient was produced by changing the mixing ratio of the two eluents: A, 0.1 % (v/v) formic acid and B, acetonitrile. The gradient started with 5 % B with a 10-min hold, was then increased to 45 % B for 60 min, and finally increased to 95 % B for a 10-min hold, following which the mobile phase was immediately adjusted to its initial composition and held for 10 min to re-equilibrate the column. The autosampler and column oven were maintained at 4 °C and 40 °C, respectively. The separated peptides were detected on the MS with a full-scan range of 300–2000 m/z (resolution of 240,000) in the positive mode followed by data-dependent MS/MS scans (resolution of 15,000). The method was set to automatically analyze the top 20 most intense ions observed in the MS scan. The ESI voltage, dynamic exclusion, ion-transfer tube temperature, and normalized collision energy were 2 kV, 30 s, 275 °C, and 30 %, respectively. The mass spectrometry data were analyzed using Proteome Discoverer 2.5 (Thermo Fisher Scientific). The protein identification was performed using Sequest HT against the protein database of *E. coli* DH5α (accession number PRJNA429943) with a precursor mass tolerance of 10 ppm, a fragment ion mass tolerance of 0.02 Da, and strict specificity allowing for up to 2 missed cleavage. Cysteine carbamidomethylation was set as a fixed modification. L-arginine (^13^C_6_, ^15^N_4_), L-lysine (^13^C_6_, ^15^N_2_), methionine oxidation, N-terminus acetylation, and N-terminal methionine loss were set as dynamic modifications. The data were then filtered at a q-value ≤ 0.01 corresponding to a 1 % false discovery rate on a spectral level.

## Data and code availability

MS data generated in this study are available in the jPOST repository^68^ (jPOST ID JPST001809). The source data are shown in **Supplementary Information 4**. The codes used in the study are shown in **Supplementary Information 5**. The other datasets generated during the current study are available from the corresponding author.

## Acknowledgments

The authors would like to thank Kei Fujiwara for useful discussions. This work was supported by JSPS KAKENHI (grant number 19K16109 and 26830139 to W.A.), JSPS Research Fellowship for Young Scientists (grant number 22J22251 to Y.K.), JST FOREST (grant number JPMJFR204K to W.A.), Sugiyama Chemical & Industrial Laboratory (http://www.sugiyama-c-i-l.or.jp/ to W.A.), and the JGC-S Scholarship Foundation (https://www.jgcs.or.jp/en/ to W.A.). The funders did not have any role in the study design, data collection and analysis, decision to publish, or preparation of the manuscript.

## Author contributions

W.A. conceived the project. Y.K., Y.M., and W.A. designed the research; Y.K., Y.M., and W.A. acquired and analyzed the data; S.A. contributed to mass spectrometric analysis; Y.K., Y.M., and M.M. contributed to the preparation of cell extracts; M.F. contributed to developing the droplet assay; M.U. advised the research. The manuscript was prepared by Y.K., Y.M., and W.A. and edited by all coauthors.

## Additional information

### Supplementary information

Supplementary Information 1. oSD·oASD pairs used in this study

Supplementary Information 2. Strains and plasmids used in this study

Supplementary Information 3. Reaction solution constituents for CF-TXTL

Supplementary Information 4. Source data

Supplementary Information 5. Codes for U-Net deep-learning and ImageJ analysis

