## Supplementary Information 5 for "Reconstitution of ribosome self-replication outside a living cell"

**Supplementary Information 5. Codes for U-Net deep-learning and ImageJ analysis**

**Script for U-Net deep learning**

from fastai.vision import *

from fastai.callbacks import *

path = untar_data(URLs.CAMVID)

print(path.ls())

path_img = path/'images

path_lbl = path/'labels

fnames = get_image_files(path_img)

lbl_names = get_image_files(path_lbl)

img_f = fnames[0]

img = open_image(img_f)

img.show(figsize=(5, 5))

get_y_fn = lambda x: path_lbl/f'{x.stem}_P{x.suffix}'

print(get_y_fn(img_f))

mask = open_mask(get_y_fn(img_f), div=True)

mask.show(figsize=(5, 5), alpha=1)

src_size = np.array(mask.shape[1:])

src_size,img.data

src_size,mask.data

class BinaryLabelList(SegmentationLabelList):

def open(self, fn): return open_mask(fn, div=True)

class BinaryItemList(SegmentationItemList):

_label_cls = BinaryLabelList

codes = np.array(["Background","Droplet"])

size = src_size

bs = 2

src = (BinaryItemList.from_folder(path_img)

.split_by_fname_file('../valid.txt')

.label_from_func(get_y_fn, classes=codes))

tfms = get_transforms(flip_vert=True)

data = (src.transform(tfms, size=size, tfm_y=True)

.databunch(bs=bs, num_workers=0)

.normalize(imagenet_stats))

data.show_batch(rows=2, figsize=(12, 9))

def acc_camvid(input, target):

target = target.squeeze(1)

return (input.argmax(dim=1)==target).float().mean()

metrics = acc_camvid

wd = 1e-2

learn = unet_learner(data, models.resnet34, metrics=metrics, wd=wd, bottle=True, callback_fns=[CSVLogger])

learn.lr_find()

learn.recorder.plot()

lr = 1e-4

learn.fit_one_cycle(20, slice(lr), pct_start=0.3)

learn.recorder.plot_losses()

learn.save('Miyawaki-stage-1')

learn.show_results(rows=2, figsize=(8, 9))

learn.unfreeze()

learn.lr_find()

learn.recorder.plot()

lrs = slice(1e-4)

learn.fit_one_cycle(100, lrs, pct_start=0.3)

learn.recorder.plot_losses()

learn.save('Miyawaki-stage-2')

learn.show_results(rows=2, figsize=(8, 9))

prediction = path/'prediction'

files = os.listdir(prediction)

count = len(files)

for i in range(count):

img = open_image(f'{prediction}/{files[i]}')

mask_pred = learn.predict(img);

mask_pred[0].save(f'{prediction}/{files[i]}_mask.png')

**Script for ImageJ**

- **Image extraction from CZI files**

dir=getDirectory("Choose a Directory");

print(dir);

splitDir=dir + "\Split\\";

print(splitDir);

File.makeDirectory(splitDir);

list = getFileList(dir);

for (i=0; i<list.length; i++) {

if (endsWith(list[i], ".czi")){

print(i + ": " + dir+list[i]);

open(dir+list[i]);

imgName=getTitle();

baseNameEnd=indexOf(imgName, ".czi");

baseName=substring(imgName, 0, baseNameEnd);

run("Split Channels");

selectWindow("C1-" + imgName);

saveAs("Png", dir + "C1-" + baseName + ".PNG");

close();

selectWindow("C2-" + imgName);

saveAs("Png", dir + "C2-" + baseName + ".PNG");

close();

run("Close All");

}

}

- **Particle analysis**

dir=getDirectory("Choose a Directory");

print(dir);

list=getFileList(dir);

for (i=0; i<list.length; i++) {

if (endsWith(list[i], "mask.png")){

print(i + ": " + dir+list[i]);

open(dir+list[i]);

imgName=getTitle();

baseNameEnd=indexOf(imgName, ".png_mask.png");

baseName=substring(imgName, 3, baseNameEnd);

open(dir + "C1-" + baseName + ".png");

selectWindow(imgName);

setThreshold(1, 255);

redirectImageTitle = "C1-" + baseName + ".png";

run("Set Measurements...", "area standard deviation mean min & max integrated density centroid limit redirect=" + redirectImageTitle + " decimal=3");

run("Analyze Particles...", "size=0-Infinity circularity=0.00-1.00 show=Outlines display results");

selectWindow("Drawing of " + imgName);

saveAs("Png", dir + "Drawing of C2-" + baseName + ".png");

close();

selectWindow("Results");

saveAs("Results", dir + baseName + ".csv");

close();

run("Close All");

}

}
